# Growth phase estimation for abundant bacterial populations sampled longitudinally from human stool metagenomes

**DOI:** 10.1101/2022.04.23.489288

**Authors:** Joe J. Lim, Christian Diener, James Wilson, Jacob J. Valenzuela, Nitin S. Baliga, Sean M. Gibbons

## Abstract

Longitudinal sampling of the stool has yielded important insights into the ecological dynamics of the human gut microbiome. However, due to practical limitations, the most densely sampled time series from the human gut are collected at a frequency of about once per day, while the population doubling times for gut commensals are on the order of minutes-to-hours. Despite this, much of the prior work on human gut microbiome time series modeling has, implicitly or explicitly, assumed that day-to-day fluctuations in taxon abundances are related to population growth or death rates, which is likely not the case. Here, we propose an alternative model of the human gut as a flow-through ecosystem at a dynamical steady state, where population dynamics occur internally and the bacterial population sizes measured in a bolus of stool represent an endpoint of these internal dynamics. We formalize this idea as stochastic logistic growth of a population in a system held at a semi-constant flow rate. We show how this model provides a path toward estimating the growth phases of gut bacterial populations *in situ*. We validate our model predictions using an *in vitro Escherichia coli* growth experiment. Finally, we show how this method can be applied to densely-sampled human stool metagenomic time series data. Consistent with our model, stool donors with slower defecation rates tended to harbor a larger proportion of taxa in later growth phases, while faster defecation rates were associated with more taxa in earlier growth phases. We discuss how these growth phase estimates may be used to better inform metabolic modeling in flow-through ecosystems, like animal guts or industrial bioreactors.

## INTRODUCTION

The human gut is an anaerobic flow-through bioreactor, ecologically distinct to each individual, that transforms dietary and host substrates into bioactive molecules important to host health ^1–3^. Disruptions to the ecological composition of the gut have been shown to mediate the progression of various complex diseases ^4–8^. Furthermore, the ecological dynamics of the gut appear to be relevant to both health and disease states ^9, 10^. However, the biological interpretation of densely-sampled adult human fecal microbiome time series is fraught.

Various dynamical models have been applied to gut microbial abundance data collected from adult human donors ^11–15^. These models often assume, either explicitly or implicitly, that day-to-day changes in abundance are proportional to population growth and/or death ^16^. However, the underlying data often do not match this assumption ^11, 16–20^. The gut is a flow-through ecosystem and commensal gut bacteria must grow fast enough to avoid dilution-to-extinction. As such, gut bacterial doubling times tend to be fast, ranging from minutes-to-hours ^21–23^. However, stool sampling frequency is usually limited to, at most, about once per day. Consequently, rapid internal population dynamics likely cannot be directly estimated from the day-to-day measurements obtained from stool ^16^.

Given these sampling limitations, and in the absence of major perturbations that require multi-day recovery processes in the human gut, it is unclear whether or not meaningful insights into commensal population dynamics can be gleaned from adult human gut microbiome time series. One workaround for inferring growth rates of bacterial populations *in situ* is to leverage metagenome-inferred replication rates ^21, 22^. Briefly, instantaneous replication rates can be estimated for abundant bacterial populations in metagenomic samples by taking advantage of the fact that fast-growing taxa show an asymmetry in reads mapping to different genomic loci, with higher read depth near the origin of replication and a lower depth near the terminus due to the initiation of multiple replication forks ^21–23^. However, even when replication rates and population abundances can both be estimated from the same metagenomic samples, it is unclear how these measurements are related to the *in situ* growth phase of a population. As such, biological interpretations regarding population size and replication rate fluctuations in flow- through ecosystems like the human gut, where internal dynamics are much faster than sampling rates, remain challenging.

Early experiments by Jaques Monod ^24^ identified distinct growth phases for bacterial populations in culture, which can be captured by the stochastic logistic growth equation (sLGE)^25^. The sLGE has been shown to be a good fit for bacterial population growth *in vitro* and in real-world, steady-state ecosystems ^26–32^. We used the sLGE to study statistical relationships between population sizes and growth rates across the various phases of growth (i.e., acceleration, mid-log, deceleration, stationary phases) to see if we could extract *in situ* growth phase information. Overall, the sLGE model yields statistical relationships that may be leveraged to identify the *in situ* growth phase of a bacterial population sampled at a regular period from a flow-through ecosystem, like the human gut.

To assess our model predictions, we sampled *Eschericial coli* populations at different points along the growth curve. We calculated population sizes and replication rates for these samples and observed excellent agreement between this *in vitro* model and our sLGE predictions. We also measured population abundance and replication rate trajectories from more than a dozen organisms across four densely sampled human gut metagenomic time series ^33^. On average, gut commensal growth rates and population sizes were positively correlated, both cross-sectionally over 84 stool donors and longitudinally within each of four stool donor time series, which suggests that most abundant taxa in the gut are growing exponentially when sampled in stool. Furthermore, we were able to identify specific growth phase signatures in abundant bacterial populations in the guts of four individuals with long and dense metagenomic time series by analyzing paired replication rate and abundance trajectories. We describe how our growth phase inference approach can serve to improve statistical inferences derived from microbiome data and to inform more accurate mechanistic modeling of flow-through ecosystems (e.g., community-scale metabolic models, which usually assume exponential growth), which could have broad implications for human health ^8, 34, 35^, agricultural systems ^36, 37^, climate change ^36, 38, 39^, and industrial bioreactor production processes ^40, 41^.

## RESULTS

### Framing the gut as an anaerobic flow-through bioreactor

The mammalian gut can be understood as an anaerobic batch culture reactor with a semi-continuous input (i.e., discrete boluses of dietary inputs, mixed with host substrates like mucin and bile acids) and output (i.e., discrete boluses of stool) ^42^, and microbial taxa must grow fast enough within the system to avoid dilution-to-extinction (**Fig. 1A**). Thus, stool sampling captures the endpoint of internal gut bacterial population dynamics. For example, in our conceptual figure we see that Taxon 1 starts growing higher up in the colon and is in stationary phase by the time a stool sample is collected, while Taxon 3 starts growing lower in the colon and is still growing exponentially at the point of stool sampling (**Fig. 1A**). Overall, the daily abundances of Taxa 1-3 represent the average (μ) steady-state population size, plus or minus some amount of biological and technical noise, at the time of stool sampling (**Fig. 1A**). To investigate improved methods for interpreting the dynamics of human gut microbial time series, we downloaded shotgun metagenomic time series data from the BIO-ML cohort (i.e., health-screened stool donors who provided fecal-transplant material to the stool bank OpenBiome) ^33^. The BIO-ML cohort contained 84 donors ^33^. To filter for dense longitudinal data, we selected a subset of donors with more than 50 time points. Four donors (i.e. donors ae, am, an, and ao) met this criterion, with 3- 5 fecal samples per week for >50 days (**Fig. 1B**).

**Figure 1.**
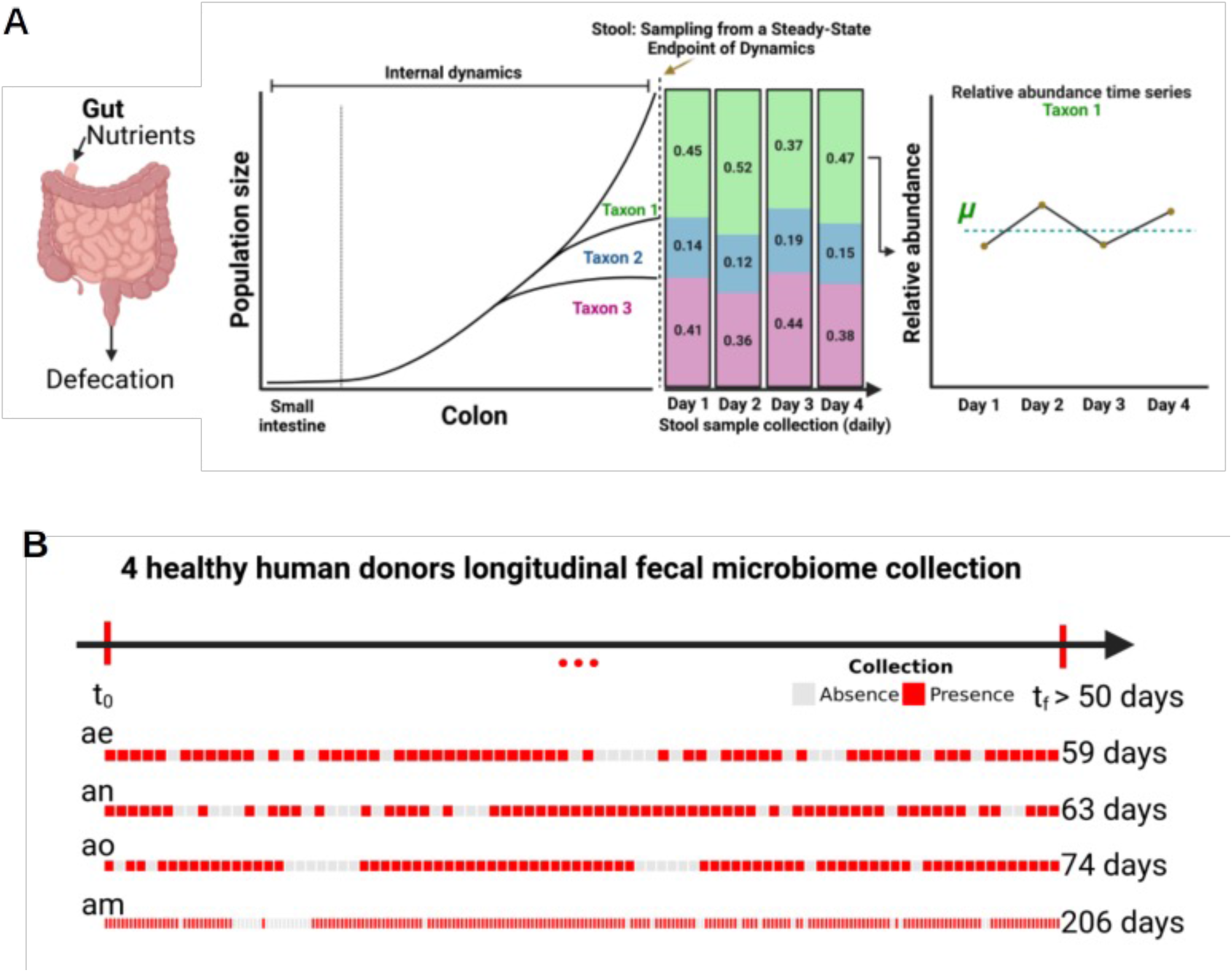
Conceptual figure showing two flow-through microbial ecosystems: a bioreactor and a human gut. **A.** Both bioreactors and guts are continuous flow-through systems. Prior to reaching the measured abundances in stool, taxa grow in the large intestine with varying growth rates, carrying capacities, and steady-state population sizes, which may be in different growth phases at the time of measurement. For example, see dynamics for Taxa 1-3 Daily stool collections show variation in abundances, but this variation likely does not reflect internal growth dynamics in the gut. **B.** Healthy BIO-ML stool donors (subject IDs: ae, am, an, and ao) with samples collected 3-5 days per week for a total of >50 time points. Red indicates presence of shotgun metagenomic sequencing data and gray represents absence of metagenomic data from consecutive daily time points.

### Characterizing the relationships between gut commensal population size and growth rate using metagenomic time series data

We first investigated the statistical properties of day-to-day fluctuations in gut bacterial population sizes, estimated from fecal shotgun metagenomic time series. Specifically, we looked at the associations between population abundance estimates (*t_n_)* and the changes in abundance estimates (i.e., deltas) between time points (*t_n+1_* – *t_n_*). Naïvely, if most bacterial populations in stool were growing exponentially, we would expect that population abundances and growth rates would be positively correlated. However, prior work has indicated an overall negative correlation between abundances and changes in abundances in stool 16S rRNA gene amplicon sequencing data generated from densely sampled human stool time series ^15^. Indeed, we found that abundant bacterial populations in the stool of the four BIO-ML donors maintained stable average abundances over time (μ), with day-to-day fluctuations above and below this average, as pictured in the example of *Bacteroides cellulosilyticus* in donor am (**Fig. 2A-B**). This kind of pattern mirrors what one would expect when randomly sampling from a stationary distribution (**Fig. 2B**). We observed that the deltas (*t_n+1_* – *t_n_*) for the same gut taxon (*Bacteroides uniformis*) measured across each donor time series, when plotted against their respective normalized abundances (*t_n_*), showed the expected negative association (**Fig. 2C**). Furthermore, similar negative associations were uniformly observed across all taxa analyzed, across all four donors (**Fig. 2D**). This negative association between population abundances and changes in abundance between time points is strongly consistent with sampling from a stationary distribution, which is equivalent to ‘regression-to-the-mean’ as an organism fluctuates around a fixed carrying capacity, similar to what we have reported previously ^15, 32^.

**Figure 2.**
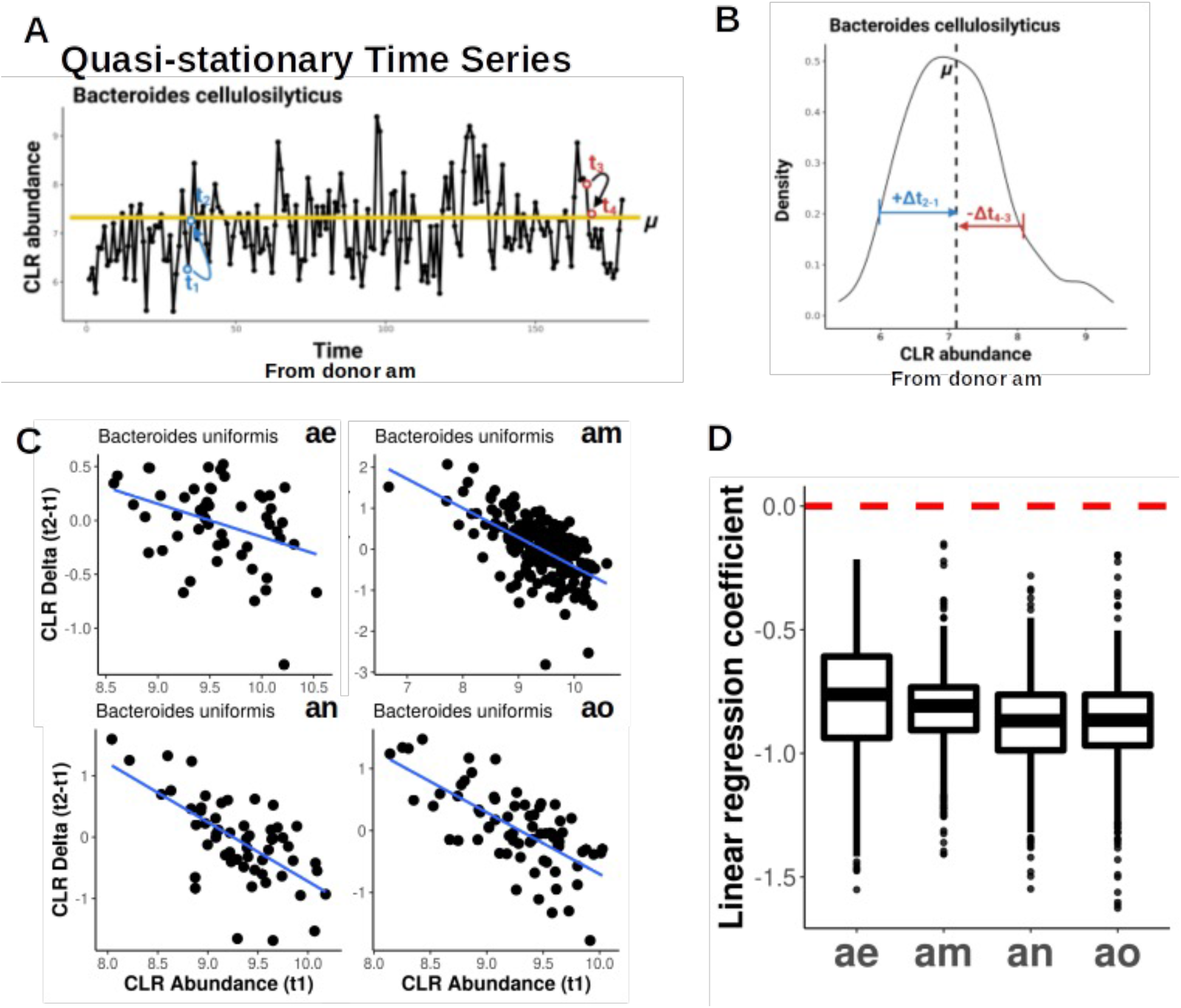
Regression-to-the-mean effect in human microbial time series data. **A.** Yellow line represents the mean abundance (μ) of *Bacteroides cellulosilyticus* over time in donor am. Time points *t_1_* and *t_3_* indicate fluctuations below and above the mean abundance, and *t_2_* and *t_4_* show the return to the mean abundance. **B.** Distribution of time series delta values (e.g., *t_2_-t_1_*) for *Bacteroides cellulosilyticus* in donor am, which is approximately normally distributed. **C.** Delta vs. abundance for *Bacteroides uniformis* time series from donors ae, am, an, and ao. **D.** Boxplots (showing median and interquartile range) of linear regression coefficients for deltas vs. abundances across all taxa time series in all four donors. Red line indicates a regression coefficient of 0.

One important ecological factor that can impact gut microbial dynamics is host diet ^43, 44^. Although changes in dietary intake can alter microbial abundance, average dietary choices are highly conserved within an individual and these choices are notoriously difficult to modify outside of radical changes in geography or lifestyle ^45–47^. Prior work demonstrated that macronutrient intake within an individual is largely stable over time and does not show significant autocorrelation or drift ^15, 48^. Indeed, for donor A from this prior study, we found that longitudinal measurements of macronutrients (i.e. daily intake of calories, carbohydrates, protein, fat, fiber, cholesterol, saturated fat, sugar, sodium, calcium) were stationary over several months, despite day-to-day fluctuations (**Fig. S1**). Combined with the overwhelming stationarity of microbial abundance trajectories within healthy individuals not undergoing major lifestyle changes ^15, 32, 33^, these results support our assertion that dietary patterns are largely stable over weeks-to-months and stool samples provide stable, steady-state population abundance estimates of abundant gut commensal bacteria.

Next, we looked at the statistical associations between calculated peak-to-trough ratios (i.e., PTRs; a proxy for growth-rate) for abundant bacterial populations from each metagenomic sample and their respective metagenomic population abundance estimates ^22^. If the deltas, presented above, were truly proportional to growth and/or death rates, we would expect that the statistical relationships between deltas and population size would be similar to those between PTRs and population size. However, unlike the regression-to-the-mean signature identified for the deltas, we found variable statistical relationships between log_2_PTR and centered log-ratio (CLR) transformed population abundances for the same taxon across the four donors (*Bacteroides ovatus*, **Fig. 3A**). Similarly, we saw a wide range of positive, negative, and null associations between log_2_PTRs and CLR abundances across all measured taxa within each donor (**Fig. 3B**). These results are inconsistent with a regression-to-the-mean signal, and suggest a more complex relationship between growth rate and population size ^49–51^. Finally, we calculated temporally-averaged (i.e., mean for all collection time within a taxon) PTRs and population sizes for each abundant taxon within each of the four donors. Overall, there was a significantly positive association (linear regression, *p-values* = 0.0318, 0.125, 0.155, 0.031 for donors ae, am, an, and ao, respectively; combined *p-value* using Fisher’s method = 0.005) between average log_2_PTR and average CLR abundance across all four donors (**Fig. 3C**), indicating that taxa with higher average population sizes tend to have higher average growth rates. This result is consistent with what one would expect to observe in exponentially-growing populations. We also looked into whether or not log_2_PTR magnitudes were inter-comparable across taxa (**Fig. S2**). We calculated log_2_PTRs for all abundant taxa detected across all 84 BIO- ML donors and found that the median log_2_PTR was fairly similar across taxonomic classes (∼0.45-0.75), with most classes showing a wide range of log_2_PTRs (**Fig. S2**). To assess whether or not log_2_PTR-CLR associations were robust to controlling for taxonomy, we included either class- or species-level categorizations as covariates in a linear regression model and saw a significant association, independent of taxonomy (class-level L = 0.0612, *p* = 8.359e ; species-level L = 0.0101, *p* = 0.0006).

**Figure 3.**
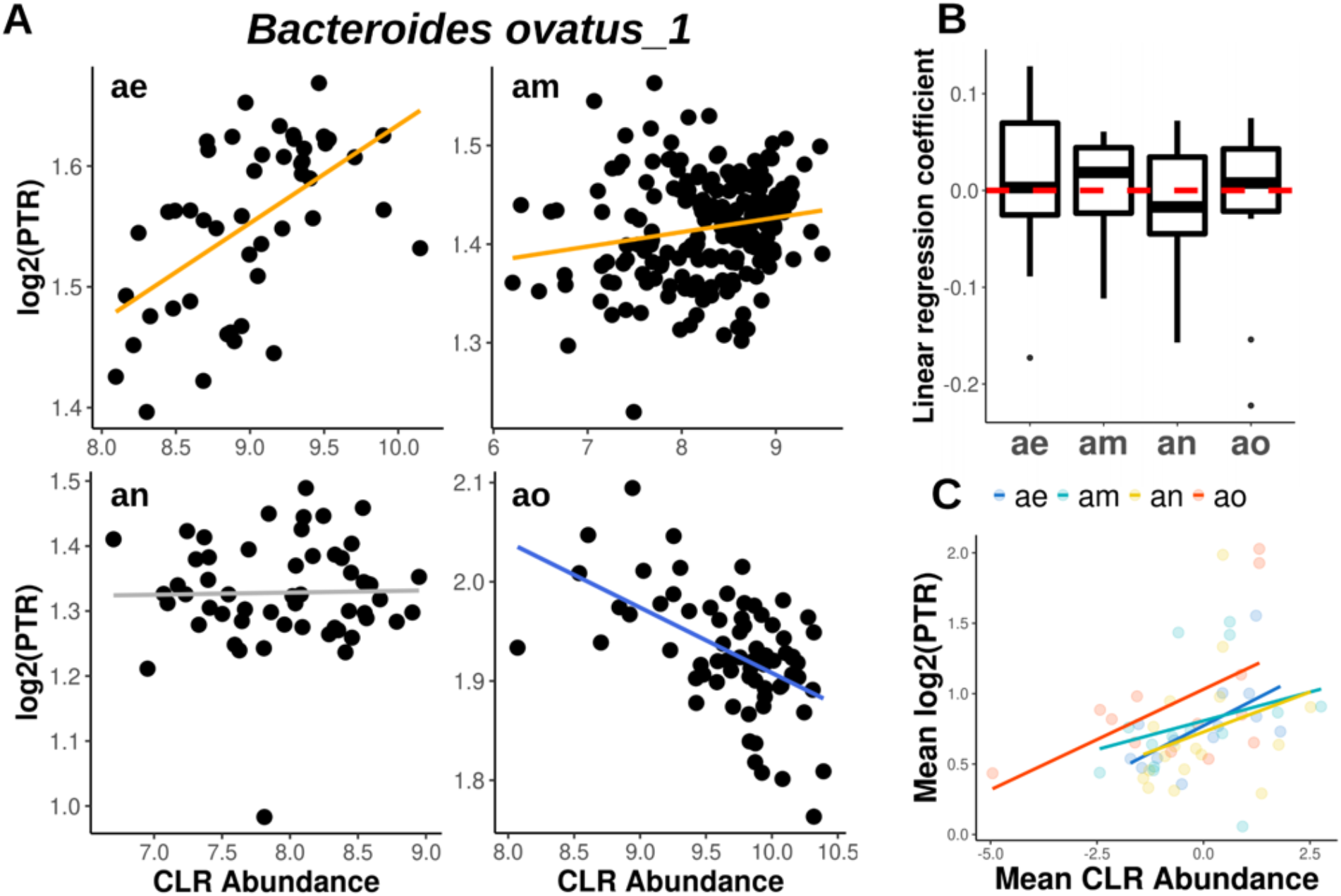
Variable relationships between PTRs and CLR-normalized abundances across human gut microbial time series. The ratio of sequencing coverage near the replication origin to the replication terminus for each species (i.e. peak-to-trough ratio, or PTR), was calculated using COPTR. **A.** Log_2_(PTR) and CLR-normalized abundance relationships for donors ae, am, an, and ao. Orange and blue lines show significantly positive and negative linear regression coefficients (linear regression, FDR adjusted *p*-value < 0.05), respectively. Gray lines indicate no statistically significant association. **B.** Boxplots (showing median and interquartile range) of linear regression coefficient combined for all filtered taxa for each donor. **C.** Mean log_2_(PTR) and mean CLR-normalized abundance for all abundant taxa in each donor (*p-values* for regressions run within each donor were combined using Fisher’s method; combined *p-value* = 0.005). PTR was calculated.

### Stochastic logistic growth equation provides insights into growth phases

In order to better understand and interpret the varying relationships we observe between log_2_PTRs and CLR abundance time series, we used a modeling approach. The basic properties of growth curves of microbial taxa can be captured using the logistic growth equation (**Fig. 4**). This model is defined such that the change in abundance for each taxon *i*(*dx_i_*/*d_t_*) is captured by the current abundance at time *t*, *x_i_*(*t*), multiplied by the maximal growth rate, *r*, and the carrying capacity (1-*x_i_*(*t*)/*k*)^52^. In this model, population size over time shows a sigmoidal curve, with the abundance asymptotically approaching *k* (Fi**g. 4A**, top panel). The derivative of this curve with respect to time yields the growth rate over time, which peaks during mid-log phase (**Fig. 4A**, middle panel). The second derivative of abundance with respect to time, which is the instantaneous change in growth with respect to time and is often referred to as the acceleration rate, shows a peak during the acceleration phase and a trough during the deceleration phase (**Fig. 4A**, bottom panel). Based on this second-derivative curve, we show the expected relationships between growth rate and abundance as you move across the logistic growth curve, along the time axis (**Fig. 4B**). These expected relationships provide a potential path forward for inferring the *in situ* growth phase of a bacterial population sampled at a consistent frequency from a flow-through ecosystem.

**Figure 4.**
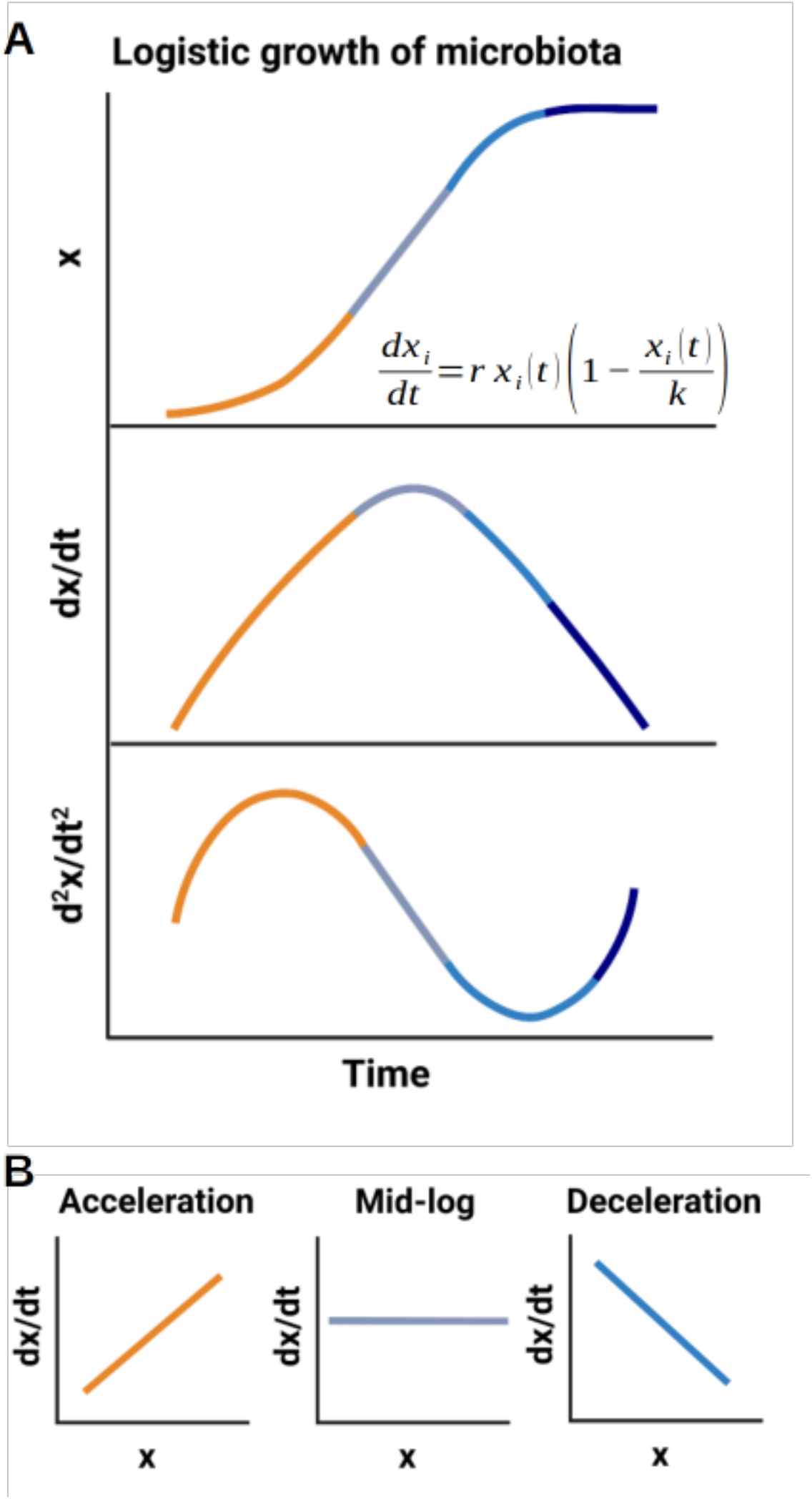
Diagram of the logistic growth equation. **A.** The logistic growth curve models abundance (*x*) with respect to time (top panel). Orange, grey, blue, and navy describes acceleration, mid-log, deceleration, and stationary phases, respectively. The first derivative of the logistic growth curve models the growth rate with respect to time (middle panel). The second derivative of the logistic growth curve models growth rate acceleration with respect to time (bottom panel). **B.** Expected relationships between abundance and growth rate at different locations along the logistic growth curve.

The logistic growth model is a deterministic equation. However, the observed abundances of commensal bacterial populations in the gut fluctuate due to myriad factors including interspecies competition, resource fluctuations, technical noise, sampling noise, and stool residence time ^27, 32, 53^. In order to approximate these fluctuations in our modeling, we introduced a stochastic term to the logistic growth model (**Fig. 5**A). Herein, *σ* denotes the noise magnitude and *ω*(*t*) represents a white noise term. Four growth phases (i.e., acceleration, mid-log, deceleration, and stationary) were defined using the half-maximum and half-minimum, respectively, of the second derivative of the LGE curve (**Fig. S3A**). We simulated 100 iterations of the stochastic logistic growth equation (sLGE) for each of a range of parameterizations (see Methods), which recapitulated the expected statistical relationships between growth rates and abundances for populations consistently sampled within our four major growth phase categories (**Fig. 5A-C**). For example, Pearson’s correlations between growth rates and abundances were significantly positive in the acceleration phase and significantly negative in the deceleration phase (**Fig. 5B**). Mid-log phase growth was more variable, but showed little-to-no significant association between growth rates and abundances (**Fig. 5B-C)**. These results were reproduced across a wide range of parameter space and were robust to varying noise levels (**Fig. S3B**).

**Figure 5.**
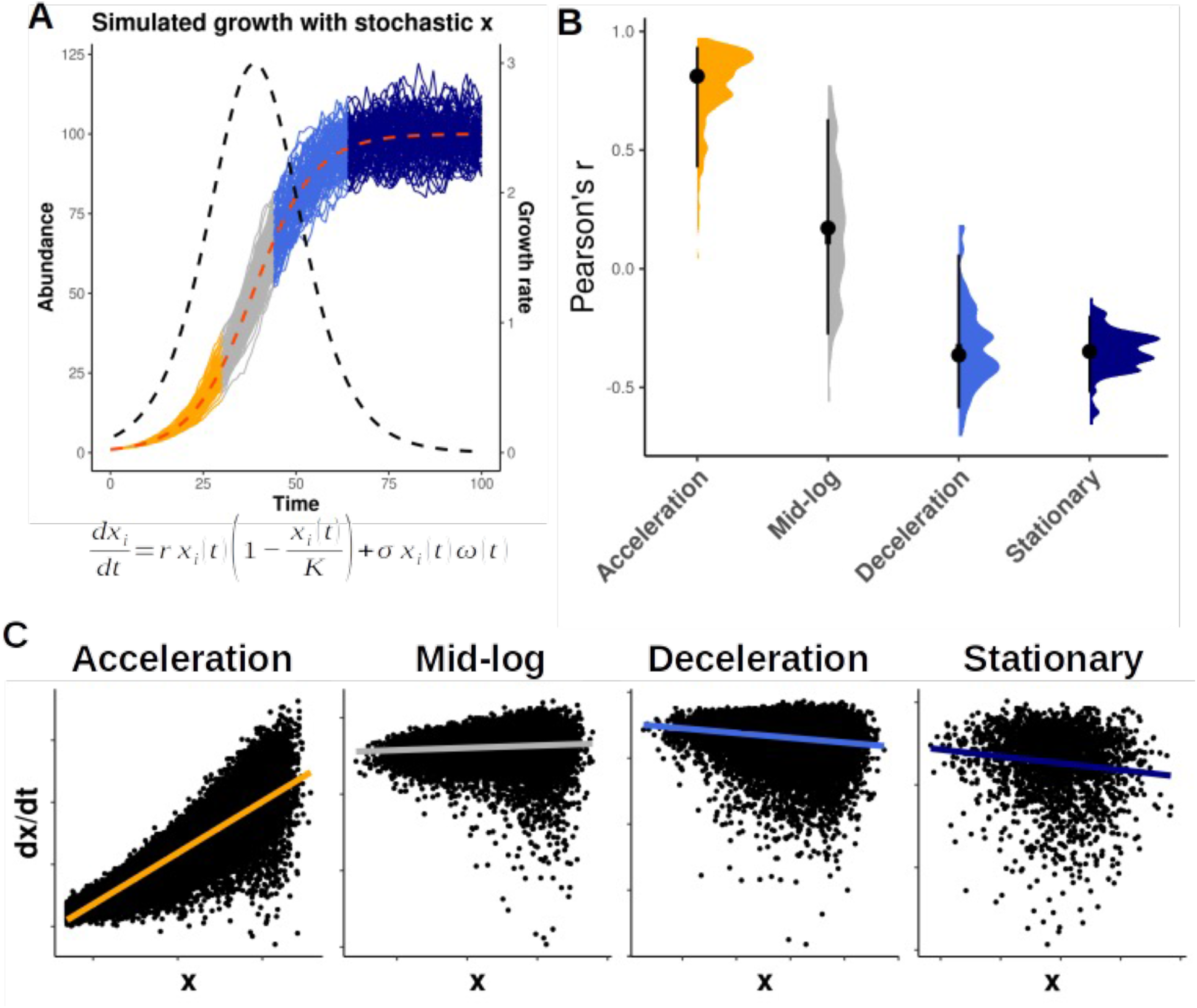
Distinguishing growth phases using the stochastic logistic growth model. **A.** Stochastic logistic growth curves with growth rate (r) = 1.2, carrying capacity (k) = 100, and noise level (n) = 0.1 across 100 iterations. Major growth phase groups in orange (acceleration), gray (mid-log), blue (deceleration), and navy (stationary). **B.** Pearson r values between abundances and growth rates in each of the four growth phase windows across variable model parameterizations (r = 1-3, k = 10-1000) and a fixed noise level (= 0.1). **C.** Scatter plots in log scale showing relationships between abundance and growth rate across the four growth phase regions defined in panel A.

Even though we expect dietary intake to be stationary within an individual, variation in diet can drive day-to-day fluctuations in the carrying capacities of microbial populations. In order to investigate whether growth-phase specific associations between abundances and growth rates were influenced by fluctuations in carrying capacity, we added variation to *k* in the sLGE model (**Fig. S4**). Fluctuations in *k* did not alter the sigmoidal shape of the sLGE curve (**Fig. S4A**), and the relationships between abundances and growth rates across growth phases were preserved (**Fig. S4B-C**).

### Validating sLGE growth phase inferences in vitro

To validate the relationship between growth rates and abundances across growth phases, we cultured replicate *E. coli* populations *in vitro* and sampled them across their growth curves (**Fig. 6A**). *E. coli* abundances were measured as OD600 values and as the log-ratio of *E. coli* reads to phiX reads (i.e., a fixed amount of the phiX genome was spiked into each DNA extraction) from the shotgun sequencing data (**Fig. 6A-C**). Growth rates were quantified as the log_2_PTR for each *E. coli* sample ^54^. The relationships between the log_2_PTRs and CLR-normalized *E. coli* abundances across growth phases matched the sLGE model predictions (**Fig. 6B-C**). Specifically, growth rates and abundances were significantly positively and negatively correlated in acceleration and deceleration phases, respectively (**Fig. 6B-C**). Furthermore, we saw no significant association between growth rates and abundances in mid-log and stationary phases (**Fig. 6B-C**). Finally, we found that samples in mid-log phase had an average log_2_PTR of 1.25±0.167 (± standard deviation), while samples in stationary phase had an average log_2_PTR of 0.358±0.059, which clearly distinguished between these phases.

**Figure 6.**
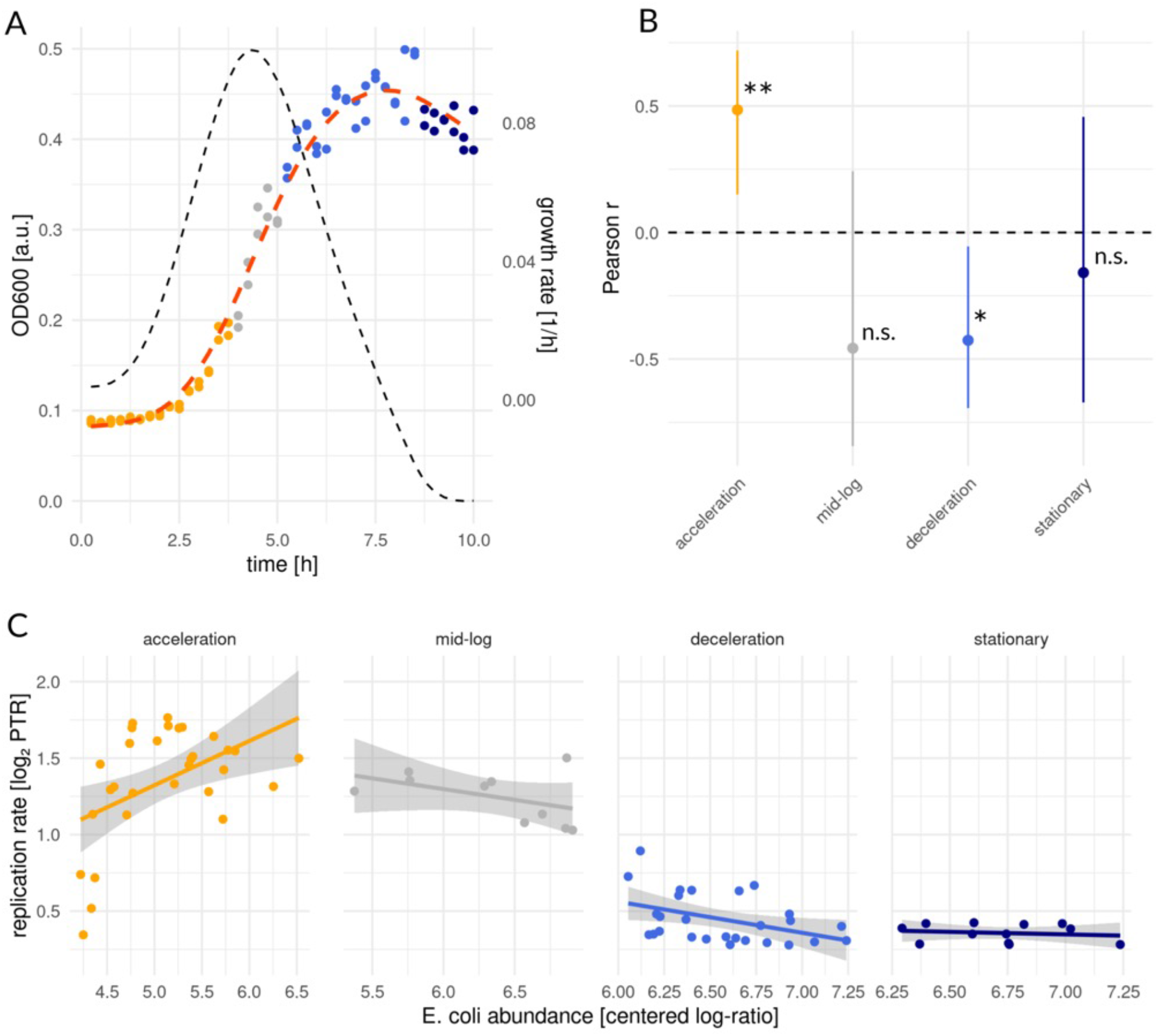
Relationship between growth rate and abundance in major growth phases in *E. coli* populations. **A.** Growth curve of *E. coli* (MG1655) using OD measurements. Colors describe major growth phases. Dotted black and red lines show the growth rate derived from OD measurements and mean growth trajectory, respectively. **B.** Pearson r values between abundance and growth rate in each of the four growth phase windows. Asterisks show statistical significance. **: p < 0.01, *: *p* < 0.05, n.s.: not significant. **C.** Scatter plots in log scale showing relationships between abundance and replication rate (log_2_PTR) across the four growth phase regions defined in panel A.

### Inferring in situ growth phases for abundant gut commensal populations sampled in metagenomic time series

Based on the sLGE results and *in vitro* validation work presented above, we assigned putative *in situ* growth phases to abundant gut bacterial populations from the four BIO-ML gut metagenomic time series. The average magnitude of the PTR provides additional information on whether a population is more likely to be in acceleration/mid-log/deceleration (i.e., log_2_PTR >> 0.358) or stationary (i.e., log_2_PTR < 0.358) phase (**Fig. 6**). For those taxa with average log_2_PTRs above the empirical stationary phase threshold, significantly positive associations (linear regression, adjusted *p*-value < 0.05, with a positive beta-coefficient) between log_2_PTRs and CLR abundances likely indicate acceleration-phase and significantly negative associations (linear regression, adjusted *p*-value < 0.05, with negative beta-coefficient) likely indicate deceleration phase. *Bacteroides cellulosilyticus*, *Bacteroides ovatus 1*, and *Megaspaera eldenii* showed significantly positive PTR-abundance associations within donor ae (**Figs. 7A and S5**). *Bacteroides xylanisolvens* had an average log_2_PTR less than the stationary threshold in donor am (**Fig. S6**). *Bacteroides ovatus* 1 and *Parabacteroides distasonis* showed positive log_2_PTR- CLR abundance associations, while *Alistipes finegoldii*, and *Bacteroides uniformis* showed negative associations in donor am (**Figs. 7A and S6**). *Acidaminococcus intestini*, *Bacteroides xylanisolvens*, and *Odoribacter splanchnicus* showed average log_2_PTR below the empirical stationary phase threshold in donor an (**Fig. S7**). *Alistipes shahii*, *Bacteroides intestinalis*, *Bacteroides thetaiotaomicron*, and *Bacteroides uniformis* showed significantly negative log_2_PTR-CLR abundance associations in donor an (**Figs. 7A and S7**). Finally, *Favonifractor plautii* showed a positive log_2_PTR-CLR abundance association and *Bacteroides fragilis*, *Bacteroides ovatus* 1, *Bacteroides uniformis*, and *Bacteroides xylanisolvens* showed negative associations in donor ao (**Fig. 7A and S8**). In all four donors, many taxa showed average log_2_PTRs greater than the stationary threshold but without significant associations between log_2_PTR and CLR abundances (**Figs. 7A and S5-8**). The absence of a significant association for these non-stationary taxa likely indicates mid-log phase, but a non-significant association could also represent a false negative (i.e., not powered enough to detect a positive or negative association with the number of time points sampled).

**Figure 7.**
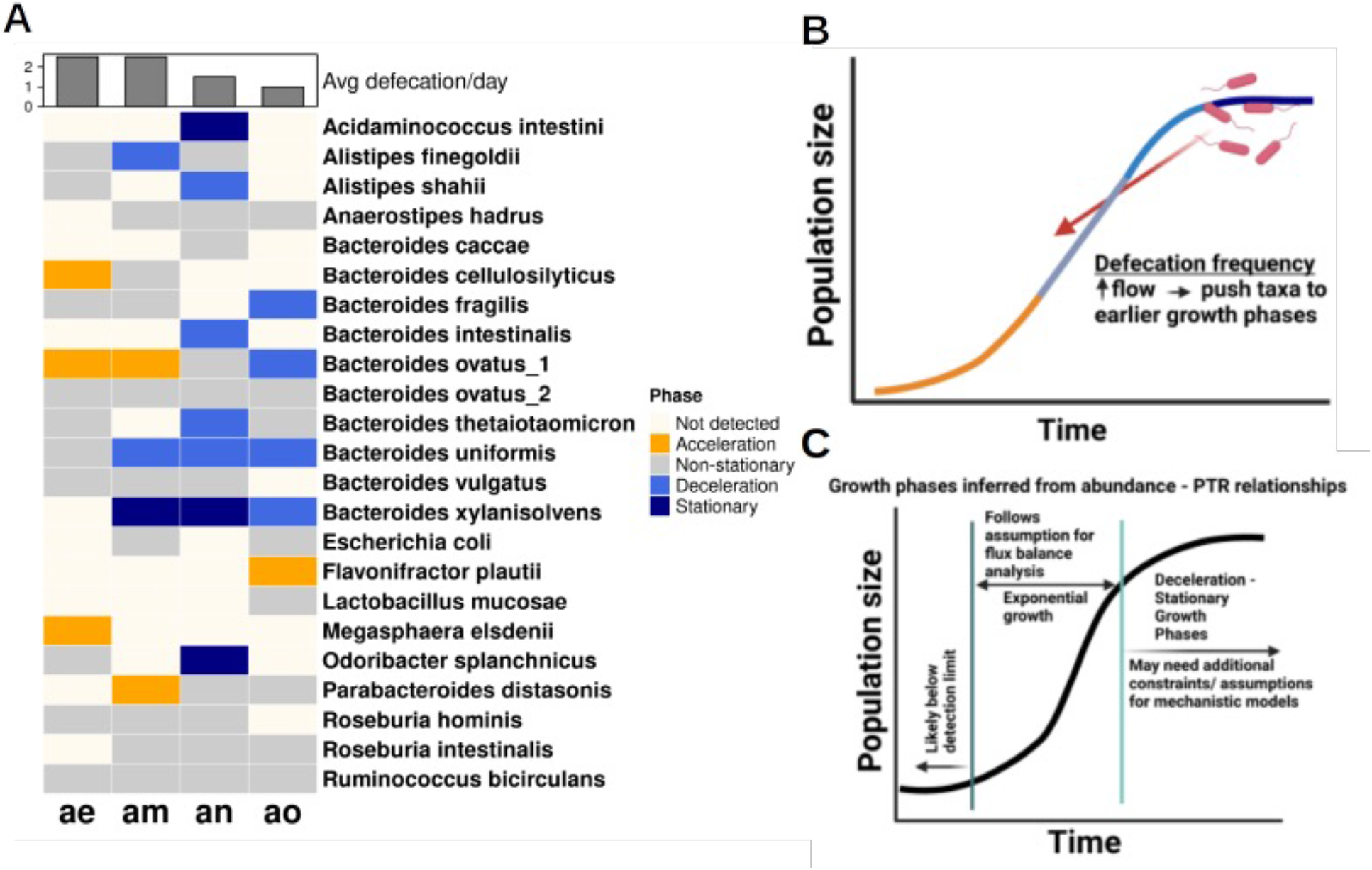
*in vivo* growth phase estimation. **A.** We find variable relationships between log_2_PTRs and population abundances across taxa in each of the four donors, consistent with the growth phase patterns observed in sLGE simulations. Donors with higher defecation rates tended to have a larger fraction taxa with positive log_2_PTR-abundance associations and fewer with negative associations, indicating acceleration and deceleration-stationary phases, respectively. Taxa in stationary phase were classified using an empirical threshold (average log_2_PTR < 0.358). Non-stationary taxa (i.e., above the stationary phase threshold, but lacking a significant correlation between log_2_PTRs and abundances) are likely in mid-log phase, but these taxa could also be in acceleration/deceleration phases (i.e., underpowered to detect the correlation). **B.** We suggest that higher defecation rates (i.e., higher dilution rates) push bacterial populations towards earlier growth phases, which is consistent with our results in panelA. **C.** Growth phase estimates can be leveraged to identify taxa that are more-or-less amenable to metabolic modeling techniques, such as Flux Balance Analysis, which assumes exponential growth.

We observed a slight difference in the number of significantly positive and negative PTR-abundance associations between donors ae/am, and an/ao. Donors ae and am tended to have a larger proportion of taxa in acceleration phase, while an and ao tended to have a larger proportion of taxa in deceleration or stationary phases. Interestingly, donors an and ao had a lower average defecation frequency (≤ 1 per day) than donors ae and am (> 1 per day). Concordantly, based on our flow-through model of the gut ecosystem (**Fig. 1A**), we would expect that bacterial populations would be pushed towards earlier growth phases at faster flow rates (**Fig. 7B**). Overall, we were able to at least partially constrain our phase estimates for all taxa with sufficient longitudinal data (**Fig. 7A**). Our approach provides a new path toward providing constraints on *in situ* growth phases for microbial populations in flow-through ecosystems.

## DISCUSSION

Many prior studies assumed, either implicitly or explicitly, that the growth and death rates of gut bacterial populations were proportional to day-to-day changes in abundances, as measured from human stool samples. However, we outline how this assumption is likely invalid due to the fact that human gut bacterial population growth/death processes inside the intestinal tract are known to be faster (minutes-to-hours) than our sampling timescales (days). In support of this assertion, we show how the statistical relationships between changes in abundance (*t_n+1_* – *t_n_*) and abundances (*t_n_*), estimated from stool metagenomic time series, indicate a regression-to-the-mean effect that one would expect when sampling from a steady-state population fluctuating around a carrying capacity (**Figs. 1-2**). Thus, as prior work has indicated ^15, 32^, bacterial taxa in the gut have stable average population sizes, which likely represent steady-state endpoints of internal dynamics (**Figs. 1-2**). Despite the fundamental mismatch between gut bacterial population dynamics and sampling timescales, we attempt to identify statistical signatures within these daily-sampled human gut time series that might provide accurate insights into *in situ* population dynamics.

While changes in abundance between time points do not appear to be related to population growth, PTRs enable direct estimates of *in situ* growth rates from metagenomic samples ^21–23, 55–57^. Unlike the relationships between deltas and abundances, which were always negative (**Fig. 2C-D**), the relationships between PTRs and abundances were quite variable (**Fig. 3A-B**). While regression-to-the-mean is a plausible mechanism for the consistent negative delta-abundance relationships (**Fig. 2**), the underlying processes driving variable log_2_PTR- abundance relationships appear to be more nuanced (**Fig. 3**).

We turned to the sLGE to explore relationships between growth rate and abundance across different phases of growth, and we found clear diagnostic patterns (**Fig. 4**). Simulations showed a wide range of demographic stochasticity (**Fig. 5**) and fluctuations in carrying capacities (**Fig. S4**) could not ablate these patterns, although adding enough noise to these models eventually overrides the signal. We validated these patterns *in vitro* and saw marked correspondence between model predictions and empirical measurements (**Fig. 5-6**). Finally, we applied our sLGE predictions to four human gut metagenomic time series. Consistent with our predictions, we found that individuals with higher defecation rates tended to be enriched for taxa in earlier growth phases (**Fig. 7**). In a recent study, we observed a similar association between PTRs and bowel movement frequency (BMF) in another independent cohort, where PTRs appeared to increase with increasing BMF ^58^. Overall, our results reveal a promising approach to inferring *in situ* growth phases for abundant organisms detected in human gut metagenomic time series.

We observed that the average log_2_PTR and average CLR abundance of a given taxon over time were positively correlated, which is consistent with exponentially-growing populations (**Fig. 3C**). However, despite this average pattern across taxa, we were also able to identify specific taxa that were abundant in stool that appeared to be in stationary phase (**Fig. 7A**). These results are highly relevant to the metabolic modeling community. Ecological interactions within free-living and host-associated microbial communities are largely governed by exchanges of small-molecule metabolites ^59, 60^. Genome-scale metabolic modeling and flux-balance analysis (FBA) has been effective mechanistic tools for simulating these metabolic exchanges, especially in controlled bioreactor systems ^61^. The objective function used to find a solution subspace for these bacterial FBA models is often biomass maximization, which assumes that these organisms are growing exponentially at steady state. Exponential growth is a valid assumption for organisms in acceleration or mid-log phases, and to some extent in deceleration phase, but this assumption breaks down completely in stationary phase. Prior work has demonstrated that biomass composition can change depending on the growth phase of a population, which ideally could be taken into account to more accurately model metabolic fluxes within the system ^62–64^. Alternatively, organisms that are not actively growing could be omitted from community-scale metabolic models of colonic metabolism ^65^. Overall, our work suggests that most abundant organisms in human stool are amenable to FBA, and our growth phase estimation approach allows for the identification of abundant populations that may not fit classical FBA assumptions.

In conclusion, we provide a new path forward for the biological interpretation of metagenomic time series data generated from adult human stool samples. Our results are somewhat reassuring for cross-sectional studies, as they indicate that bacterial abundances in the gut fluctuate around stable carrying capacities within an individual, making inter-individual comparisons fairly robust. Furthermore, this suggests that multi-day averages of abundances will be even more accurate estimates of this carrying capacity, as we have suggested previously^33^. This work is especially relevant to the design and interpretation of human gut microbiome studies that aim to characterize or investigate ecosystem-scale dynamics. We hope that *in situ* growth phase estimation will be applied more broadly to other kinds of flow-through environments to improve our understanding of internal dynamics in these systems and provide improved constraints for mechanistic modeling of microbial communities.

## METHODS

### Stationarity testing for daily nutrient intake in a human stool donor

Metadata for daily nutrient intake, excluding the time window when the donor was traveling abroad, was downloaded from David et al. ^48^. We tested for stationarity in these nutrient intake time series using the augmented Dickey-Fuller (ADF) test (tseries package in R ^66^), with significance threshold for stationarity at *p* < 0.1. ADF tests the null hypothesis that a unit root is present in a time series, with the alternative hypothesis being that the time series is stationary. Thus, significant p-values indicate stationarity of the time series. All analyses throughout the manuscript in R were conducted in R v4.2.2 ^67^, unless stated otherwise.

### *E. coli* strain information and growth curve analysis with a microplate reader

*Escherichia coli* strain (MG1655) was streaked from a glycerol stock onto R2A agar plates (Thermo Fisher Scientific: Oxoid CM0906) and incubated overnight at 37°C. A colony was selected using an inoculating loop and transferred to 200 mL of LB-broth (Lennox) and grown at 37°C overnight in a shaking incubator at 225 rpm until the culture reached stationary phase. The overnight culture was then diluted in fresh LB medium to an OD of 0.51 (600 nm). The diluted culture was then chilled for ∼25 minutes at ∼2°C using an ice bath to synchronize metabolic activity. The chilled culture was then aliquoted (2μL) into a non-treated 96-well flat-bottomed plate (Thomas Scientific Cat No. 1154Q44) containing 198 μL of LB media (Lennox) in each well. The inoculated plate was then transferred to a BioTek Epoch II plate reader set to 37°C with orbital shaking and programmed to make OD600 readings every minute for the first 60 minutes and every 5 minutes for the remainder of the experiment (∼10 hours). The first set of inoculations covered plate rows A and B (n = 24), this was followed by the sequential inoculation of the next 3 sets of rows at 15-minute intervals (i.e., Set 1 = A/B: 0 min; Set 2 = C/D 15 min; Set 3 = E/F: 30 min; Set 4 = G/H 45 min.). This resulted in 4 sets of replicate cultures inoculated 15 minutes apart, allowing sampling every hour for the next 10 hours, spanning 40 time points spaced 15 minutes apart. To ensure there was enough DNA for sequencing at early low OD time-points (first two sample points), we pooled two wells into one sample. All samples were collected in PCR strip tubes (Axygen: PCR-0208-CP-C) and centrifuged at room temperature to pellet the cells. The supernatant was decanted and the remaining cell pellet was immediately frozen in liquid nitrogen for storage at -80°C.

### DNA extraction, library preparation, and sequencing

Cell Pellets were resuspended and transferred to 96 deep-well plates for DNA extraction using the IBI Scientific 96-well Genomic DNA Bacteria Kit (IBI Scientific: IB47295) per the manufacturer’s protocol. DNA quantification was done using Qubit HS DNA assay, on a Qubit3 device. After DNA quantification, we added PhiX DNA (Thermo Fisher Scientific: SD0031) as an internal standard and run-quality monitor across all samples. A total of 500 fg PhiX DNA was added to each DNA sample before library preparation. DNA libraries were constructed following the NEBNext Ultra II FS DNA Library Prep Kit for Illumina (New England Biolabs: E7805L) and indexed using Dual Index Primer Set 2 (New England Biolabs: E7780S). Libraries were quantified again via Qubit 3, and the quality and size of libraries were checked using an Agilent Tapestation, and a D5000 high sensitivity DNA tape assay. Libraries were pooled to 2 nM and sent to NovoGene for sequencing on a NovaSeq 6000 device (Illumina, USA). A partial lane was used for sequencing, 150 cycles, generating ∼64GB (∼3.3 million reads per sample) of paired-end reads.

### Shotgun metagenomics data processing and analysis

Longitudinal shotgun metagenomics sequencing data from healthy human stool samples (BIO- ML) was downloaded from NCBI BioProject accession PRJNA544527, and the associated metadata was downloaded from the associated article ^33^. Raw FASTQ files from the BIO-ML cohort and from the *in vitro E. coli* experiment were filtered and trimmed using FASTP ^68^, removing the first 5 nucleotides of the read 5’ end to avoid leftover primer and adapter sequencing not removed during demultiplexing and an adaptive sliding window filter on the 3’ end of the read with a required minimum quality score of 20. Reads containing ambiguous base calls, having a mean quality score less than 20, or with a length smaller than 50nt after trimming were removed from the analysis. Taxonomic assignment on the read level was performed with Kraken2 using the Kraken2 default database ^69^. Abundances on the kingdom, phylum, genus, and species ranks were then obtained using Bracken ^70^. Trimmed and filtered reads were then aligned to 2,935 representative bacterial reference genomes taken from the IGG database (version 1.01) using Bowtie2 ^71, 72^. Coverage profiles and log_2_ estimates of peak-to-trough ratios (PTRs) were estimated using COPTR v1.1.2 at the species-level within each sample for taxa that passed our abundance threshold ^54^. PTR estimates were then merged with Bracken abundance estimates, retaining only those species identified by both methods (Kraken2 and Bowtie2 alignment to IGGdb). For the *in vitro E. coli* experiment, reads were aligned to a custom database containing the *E. coli K12* strain genome (NCBI accession NC_000913.3) and the phiX174 genome (NCBI accession NC_001422.1). CLR abundances were then calculated from the read counts for the *E. coli* genome and the phiX174 genome.

The processed data containing the raw reads and log_2_ peak-to-trough ratios (log_2_PTRs) were read into R version 4.1.3 for analysis ^67^. All plots were generated using ggplot2 ^73^, unless indicated otherwise. BIO-ML donors were selected by retaining individuals with over 50 metagenomic time points, resulting in four time series (i.e., donors ae, am, an, and ao). Distinct *Bacteroides ovatus* strains across all four donors contained duplicated taxon names with unique taxonomic identifiers, and were renamed to “*Bacteroides ovatus_1*” and “*Bacteroides ovatus_2.*” Raw read counts for a given taxon within a sample were centered log-ratio (CLR) transformed^74^. Taxa that had matched log_2_PTR and CLR abundance information available across more than 5 time points within an individual, with time differences between samples less than three days, were used in subsequent analyses. Changes in normalized abundance were calculated as *Abundance changes*(*delta*) = *x*(*t* + 1) - *x*(*t*), where Δ*t* < 3 *days*. To assess the regression-to-the-mean effect, CLR-normalized abundances were plotted against deltas for each taxon, and the regression coefficients, aggregating all microbial taxa, were plotted as boxplots (showing median and interquartile range), summarized by donor.

For each donor, to estimate the growth phase of each individual taxon, we used linear regression of CLR-normalized abundances vs. log_2_PTRs, followed by a Benjamini-Hochberg p-value correction to control for the false discovery rate (FDR) in base R. Regression coefficients with FDR-adjusted p-values < 0.05 were considered significant. Taxa with average log_2_PTRs < 0.358 (experimentally-determined stationary threshold) were designated as being in stationary phase. For those taxa not designated as being in stationary phase, significantly positive or negative associations between log_2_PTRs and abundances were considered to be in acceleration or deceleration phase, respectively. Those with no correlation and an average log_2_PTR above the stationary threshold were constrained to be in mid-log phase or in acceleration/deceleration phase (i.e., if there was a false negative due to lack of statistical power in detecting a positive or negative slope). Linear regression was also used to test whether or not average CLR-normalized abundances and average log_2_PTRs were significantly associated within each donor, and p-values from individual tests were combined using Fisher’s method ^75^.

### Stochastic logistic growth model simulation

The stochastic logistic growth equation (sLGE) was implemented as: 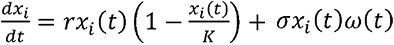, where *t* is time, *r* is the growth rate, *x_i_* is the abundance of taxon *i*, *K* is the carrying capacity, *σ* is the noise magnitude term, and *ω*(*t*) is the noise distribution term. Using the R package sde ^76^, taxonomic growth was simulated with *x_i_*,0 = 1, *t*_0_ = 1 to *t_final_* iterations. The other parameters were varied as described in the results and below. To investigate the impact of noise on sLGE trajectories, noise levels were set from 0.001 to 1, with *r* and *K* ranging from 1 to 3 and 10 to 1000, respectively. To investigate the statistical relationships between deltas and abundances across growth phases and across model parameterizations, Pearson’s R coefficients and p-values were calculated for each of the three growth phase categories. The growth phases for each model parameterization were defined using the non-stochastic logistic growth equation 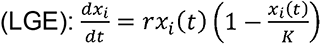, the solution for which can be written as 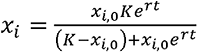.

The *x_i_* values for each simulated time point from solving the LGE were used to calculate the first derivative (i.e., the growth rate), which is exactly equal to the LGE. The second derivative (i.e., growth acceleration), 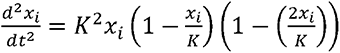, was calculated using solved *x_i_* values. Growth phases from the sLGE were defined using the second derivative curves. First, the intersections of the acceleration curve and the half-max, *a*_1_ and *a*_2_, and the half-min, *a*_3_ and *a*_4_, were calculated (**Fig. S3**). The corresponding simulated time points of *a_j_*, denoted as *s_j_*, where *j = 1 - 4*, were then used to define growth phases as follows: lag phase: *t* < *s*_1_; acceleration phase: *s*_1_<*t*<*s*_2_; log phase: *s*_2_<*t*<*s*_3_; deceleration phase:*s*_2_<*t*<*s*_4_; and stationary phase:t>*s*_4_. Here, lag and acceleration phases were combined, as these phases display similar delta-abundance relationships along the logistic growth curve. Conceptual diagrams were created using BioRender.

Death or dilution terms were not explicitly added to the simulated sLGE models. Here, we discuss how death or dilution rates are equivalent to changing the carrying capacity term, which has no impact on our growth phase inferences. Analytically, a decrease in abundance at a given time can be represented as a fraction of the current abundance subtracted from the 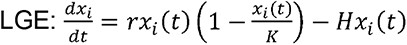. Here, *H* is the “harvest rate”, which determines the proportional decrease in each timepoint in the equation. At steady state, 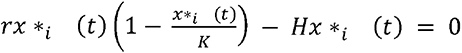, where x_\**i*_ (*t*) represents the fixed point. Two equilibria exists in this equation: x_\**i*_ (*t*) = 0 and 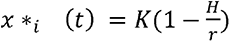, with the latter being asymptotically stable. As *H* increases, the stable population size x_\**i*_ (*t*) = 0 decreases due to the proportional decrease in *K*. As long as *H* does not exceed the intrinsic growth rate of gut microbes, which is expected for highly abundant and stably colonized taxa, the resulting *K* becomes the new stable *K*. To show that variation in *k* does not impact the relationship between growth rate and abundance, we simulated the LGE with stochastically varying *K* by adding the stochastic term, i.e. *σk_i_*(*t*)*ω*(*t*), to *k_i_*(*t*) (**Fig. S4**). In base R, simulation was performed for 100 iterations with the same noise levels (*σ* = 0.1) as the representative sLGE simulations with stochastic . Major growth phases were defined the same way as sLGE simulations with stochastic .

### Data and code availability

Nextflow pipelines implementing the processing of metagenomic shotgun sequencing data from raw reads to taxonomic abundance matrices and PTR estimates can be found at https://github.com/Gibbons-Lab/pipelines/ (metagenomics pipelines). These DNA datasets are publicly available from the National Center for Biotechnology (NCBI) Sequence Read Archive (SRA), accession code PRJNA942341. Scripts used to analyze the data, run the sLGE simulations, and produce the figures in the manuscript have been deposited at https://github.com/Gibbons-Lab/human-microbiome-time-series-growth-phase-estimation.

## Acknowledgements

We would like to thank Shijie Zhao for suggesting that we investigate PTR-abundance relationships in the BIO-ML data set. Thanks to Pamela Troisch for help with DNA library preparation for the *in vitro* experiment. We would also like to thank Nitin Baliga, Amy Willis, Julia Cui, and the members of the Gibbons Lab for helpful discussions of this work. SMG and CD were supported by a Washington Research Foundation Distinguished Investigator Award and by startup funds from the Institute for Systems Biology. JL was supported by the Environmental Pathology/Toxicology training grant (ES007032). Research reported in this publication was supported by the National Institute of Diabetes and Digestive and Kidney Diseases of the National Institutes of Health (NIH) under award number R01DK133468 (to SMG).

## Supplemental Figures

**Figure S1.**
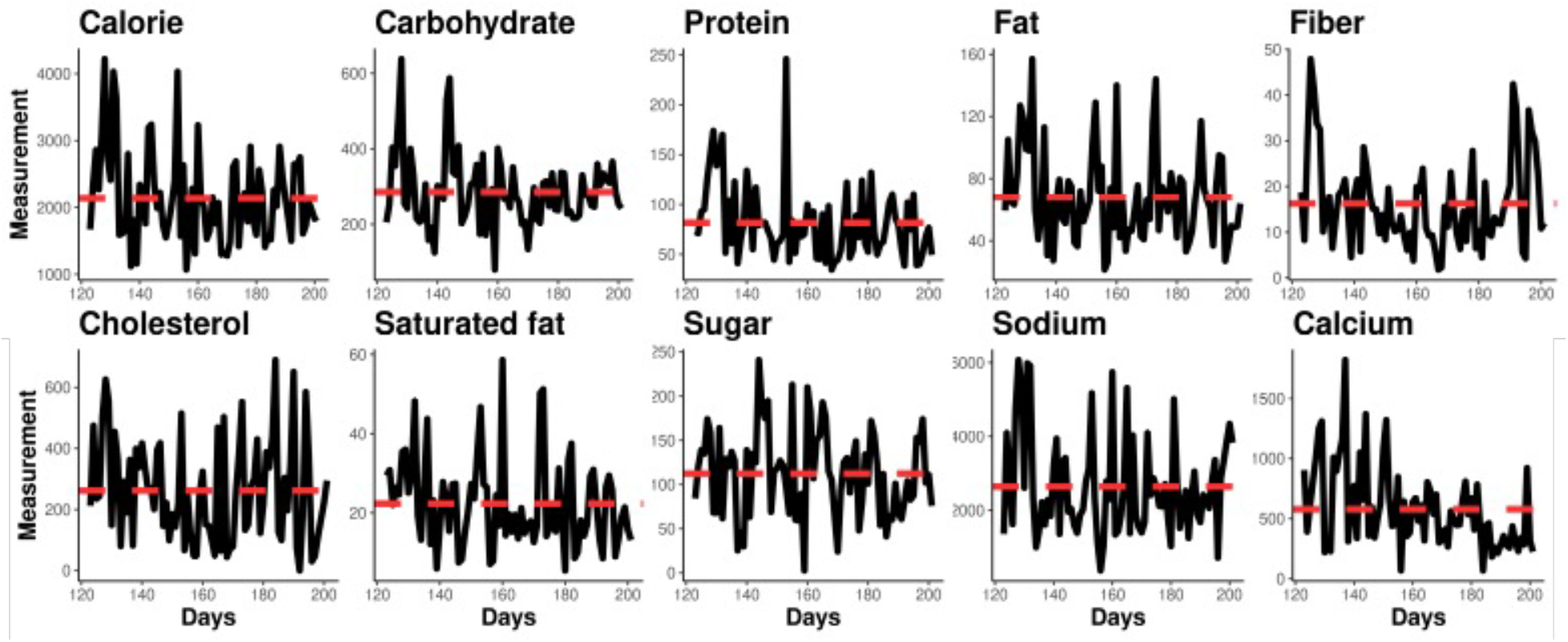
Lack of autocorrelation detected in most daily macronutrient intake. Daily measurements of nutrient intake were downloaded from David et al. ^48^. Post-travel time points are shown. Autocorrelation was tested using the augmented Dickey-Fuller test (i.e., p < 0.1 indicates significant stationarity of a dietary variable). For each nutrient, p-values are reported for the recordings post-travel period of the subject. Calorie (*p* = 0.0341), carbohydrate (*p* = 0.0144), protein (*p* = 0.0314), fat (*p* = 0.0172), fiber (*p* = 0.0369), cholesterol (*p* = 0.0534), saturated fat (*p* < 0.01), sugar (*p* = 0.0123), sodium (*p* = 0.0341), calcium (*p* < 0.01). Dotted red lines show the mean measurement values.

**Figure S2.**
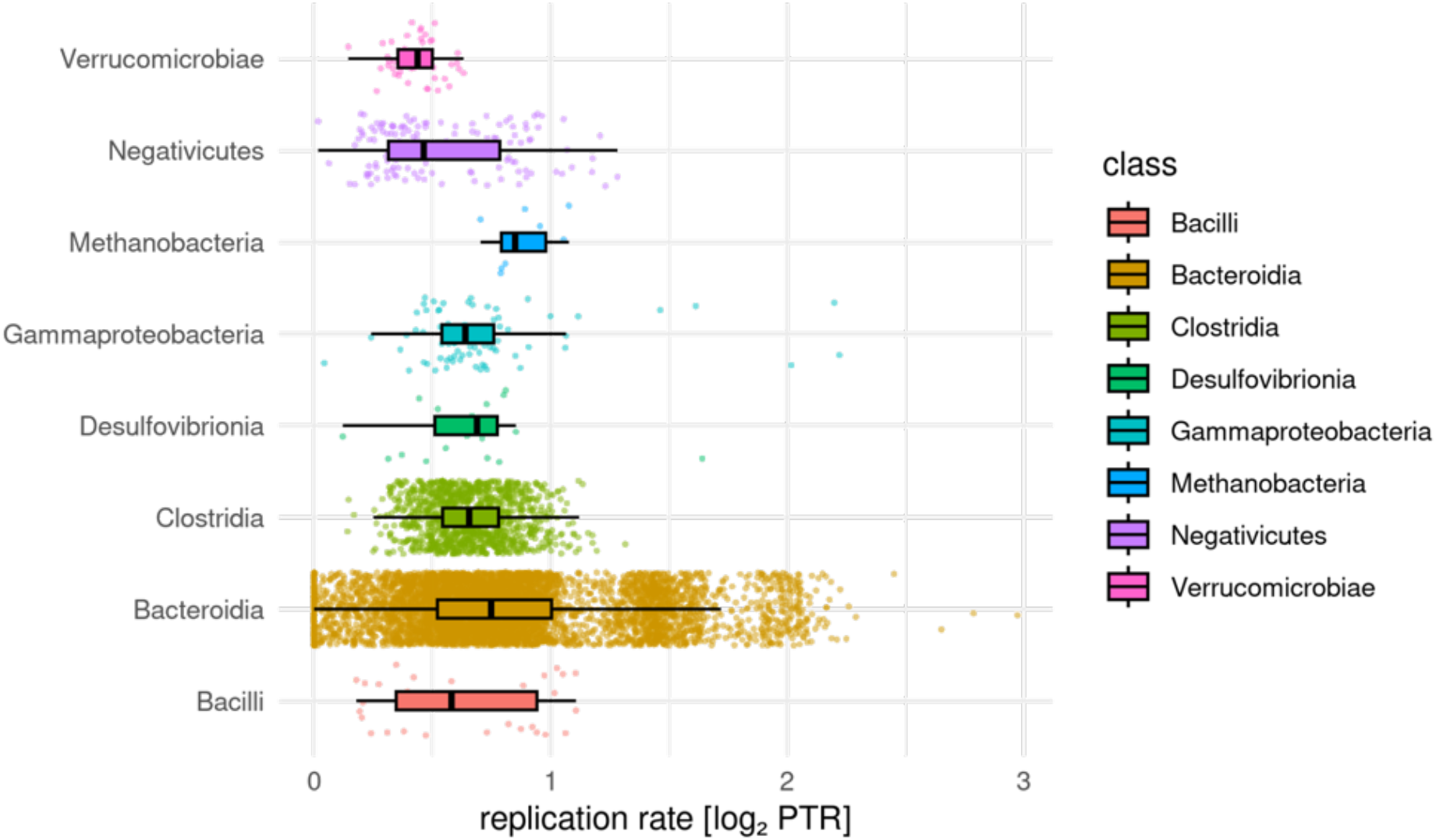
Distributions of log_2_PTR values across 84 BIO-ML donors, broken down by phylogenetic class. We see a fairly wide range of log_2_PTRs within each taxonomic class. The median log_2_PTR across classes varies between ∼0.45 and ∼0.75. In a linear regression model, controlling from taxonomic group as a covariate, we see a significant positive association between log_2_PTRs and CLR abundances at the class-level (= 0.0612, *p* = 8.359e^-60^). This positive taxonomy-controlled association is preserved at the species-level (= 0.0101, *p* = 0.0006).

**Figure S3.**
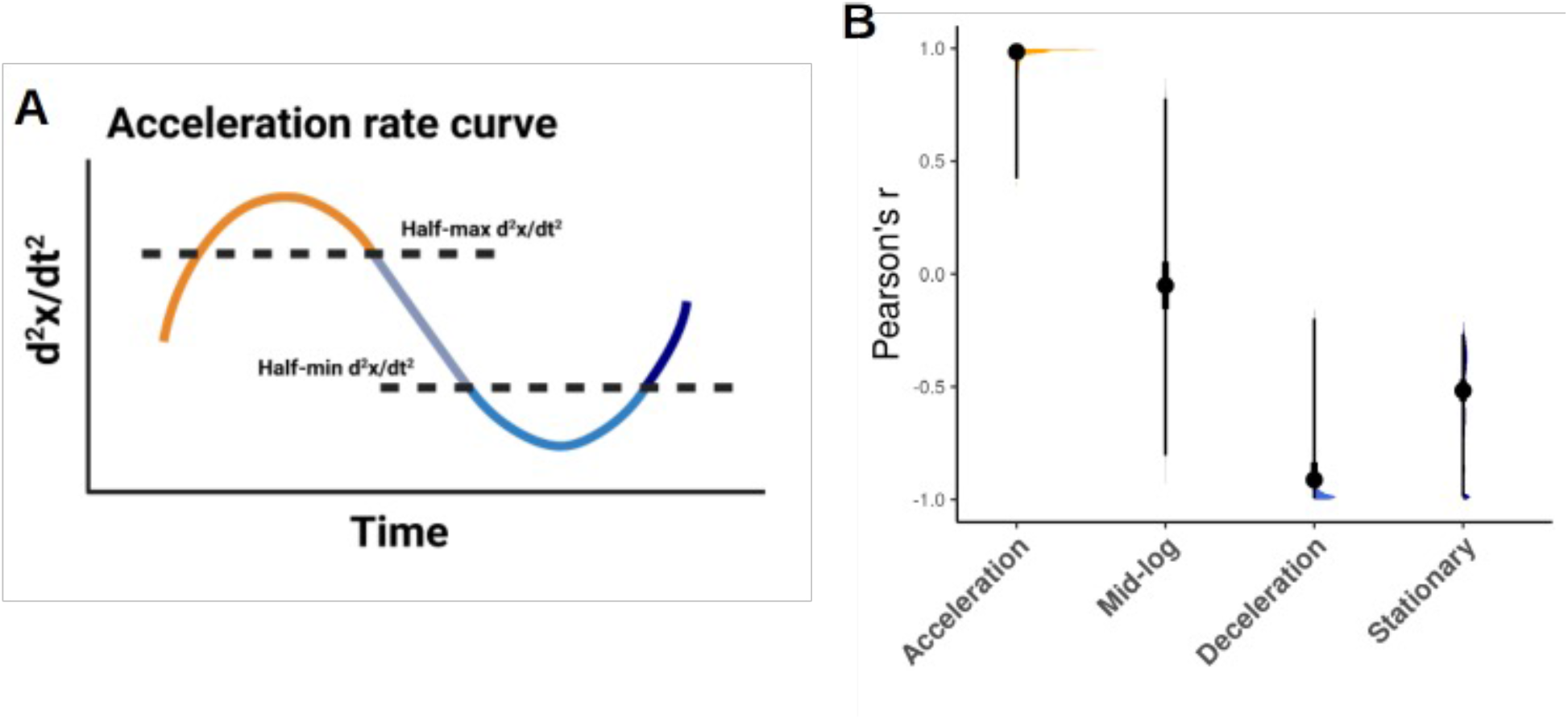
Definition of major growth phases using the stochastic logistic growth model. **A.** The half-maximum of the peak and half-minimum of the trough of the second derivative of abundance were used to define growth phases across model parameterizations. **B.** Pearson r values between abundances and growth rates in the three growth phase categories obtained from combined sLGE simulation results across a range of growth rates (r = 1-3), carrying capacities (k = 10-1000), and noise levels (n = 0.001-1).

**Figure S4.**
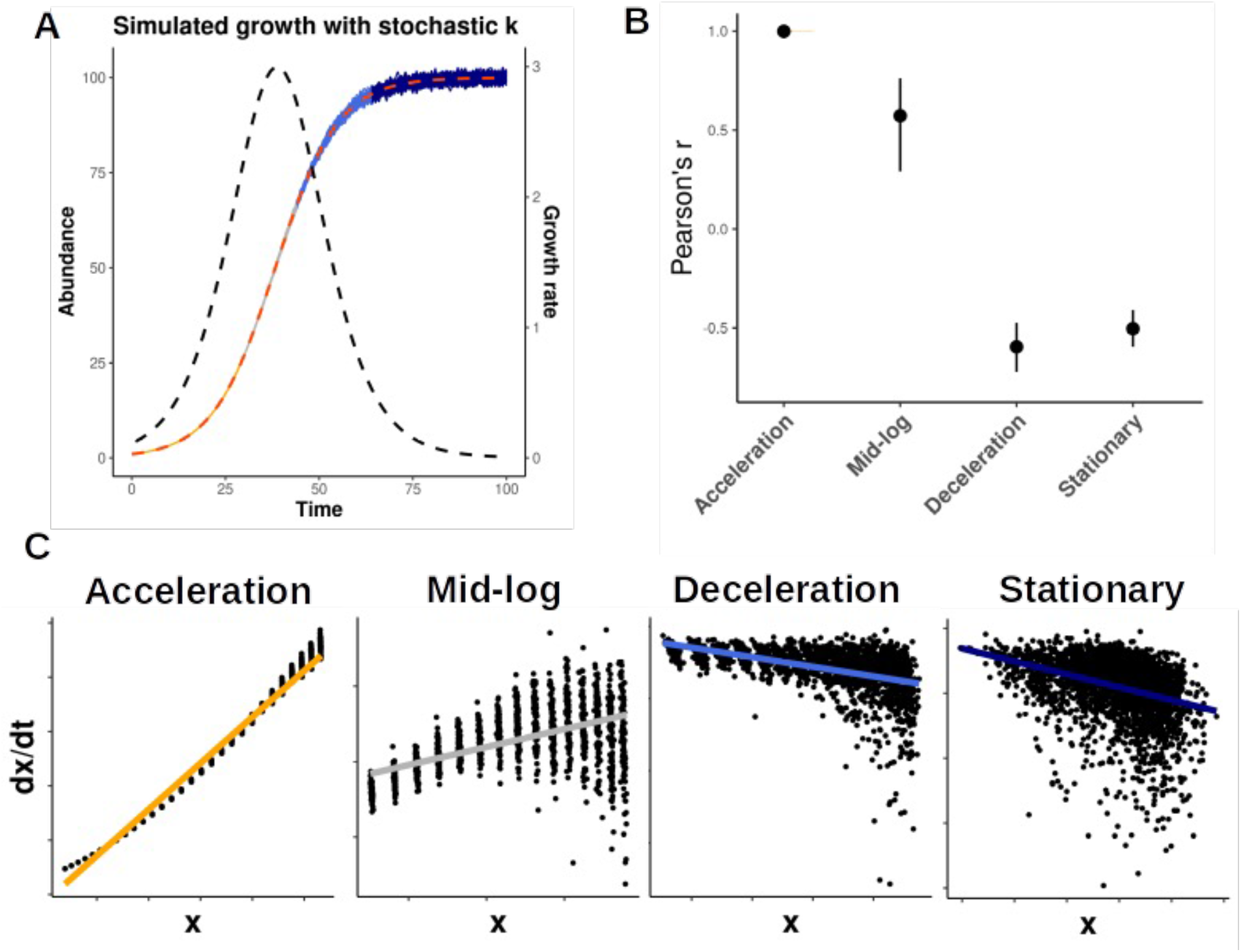
Relationship between growth rate and abundance across the major growth phases, simulated using the logistic growth model with stochastically varying carrying capacities. A. Stochastic logistic growth curves with growth rate (r) = 1.2, carrying capacity (k) = 100, and noise level (n) = 0.1 applied to k across 100 iterations. Major growth phase groups in orange (acceleration), gray (mid-log), blue (deceleration), and navy (stationary). B. Pearson r values between abundances and growth rates in each of the four growth phase windows across variable model parameterizations (r = 1-3, k = 10-1000) and a fixed noise level (= 0.1). C. Scatter plots in log scale showing relationships between abundance and growth rate across the four growth phase regions defined in panel A.

**Figure S5.**
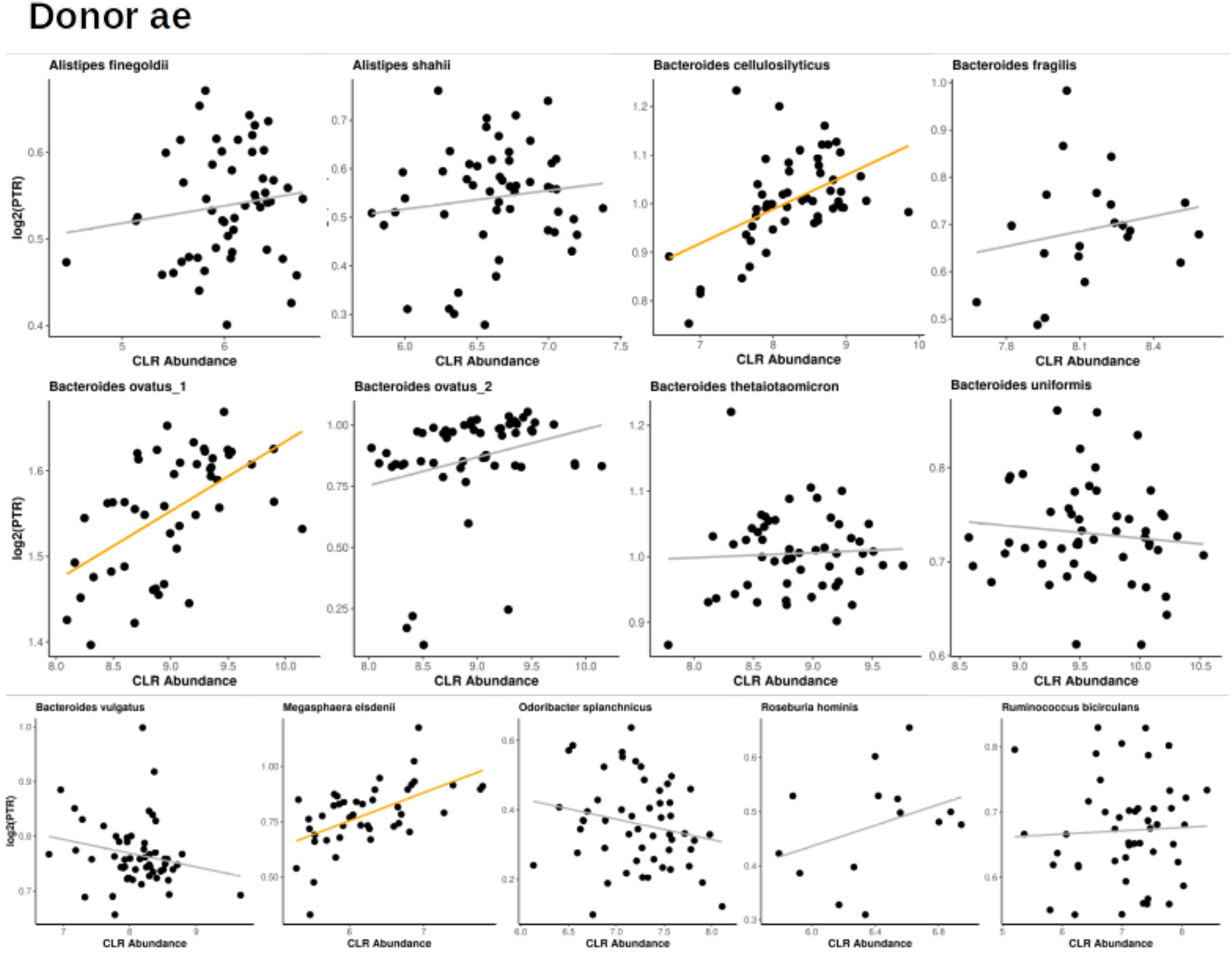
Relationships between abundance and log2(PTR) for abundant taxa in donor. **ae**. Abundant taxa with relatively dense longitudinal PTR and abundance data (at least 5 matched data points; time differences between adjacent samples less than three days) were selected for analysis. Gray trend lines show no significant correlations, orange trend lines indicate significant positive correlations, and blue trend lines represent significant negative correlations (linear regression, BH-FDR < 0.05).

**Figure S6.**
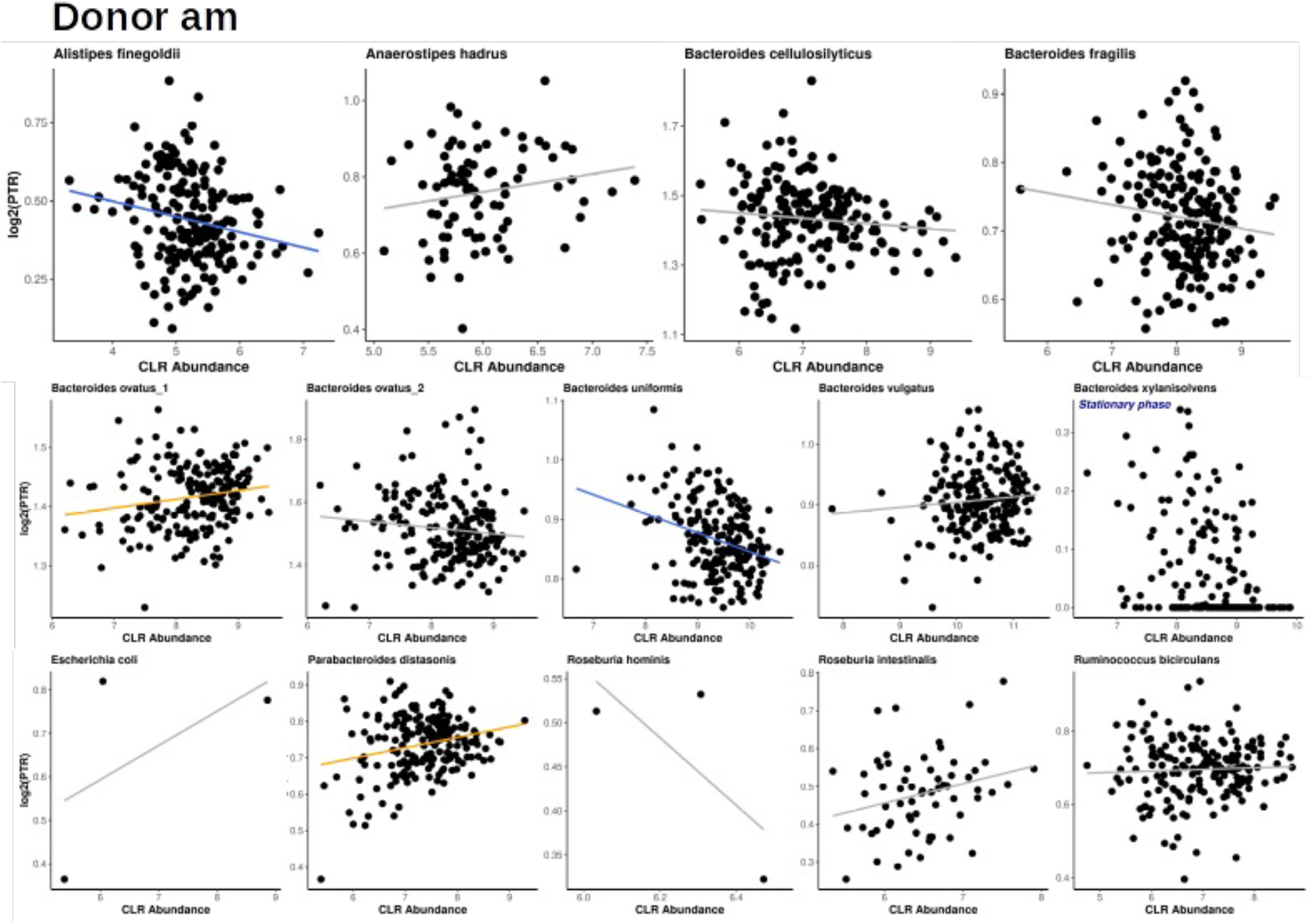
Relationships between abundance and log2(PTR) for abundant taxa in donor. **am**. Abundant taxa with relatively dense longitudinal PTR and abundance data (at least 5 matched data points; time differences between adjacent samples less than three days) were selected for analysis. Gray trend lines show no significant correlations, orange trend lines indicate significant positive correlations, and blue trend lines represent significant negative correlations (linear regression, BH-FDR < 0.05).

**Figure S7.**
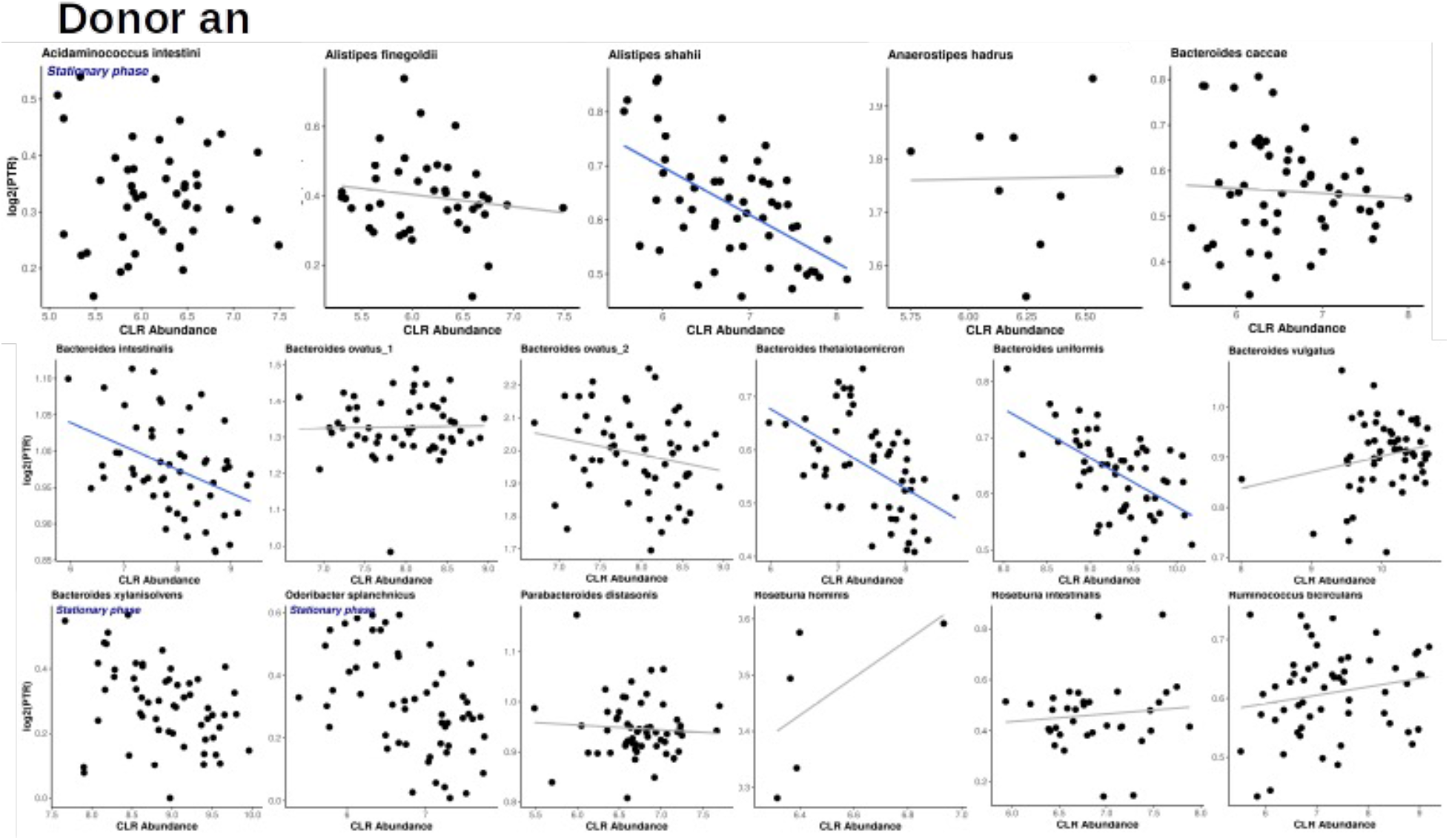
Relationships between abundance and log2(PTR) for abundant taxa in donor. **an**. Abundant taxa with relatively dense longitudinal PTR and abundance data (at least 5 matched data points; time differences between adjacent samples less than three days) were selected for analysis. Gray trend lines show no significant correlations, orange trend lines indicate significant positive correlations, and blue trend lines represent significant negative correlations (linear regression, BH-FDR < 0.05).

**Figure S8.**
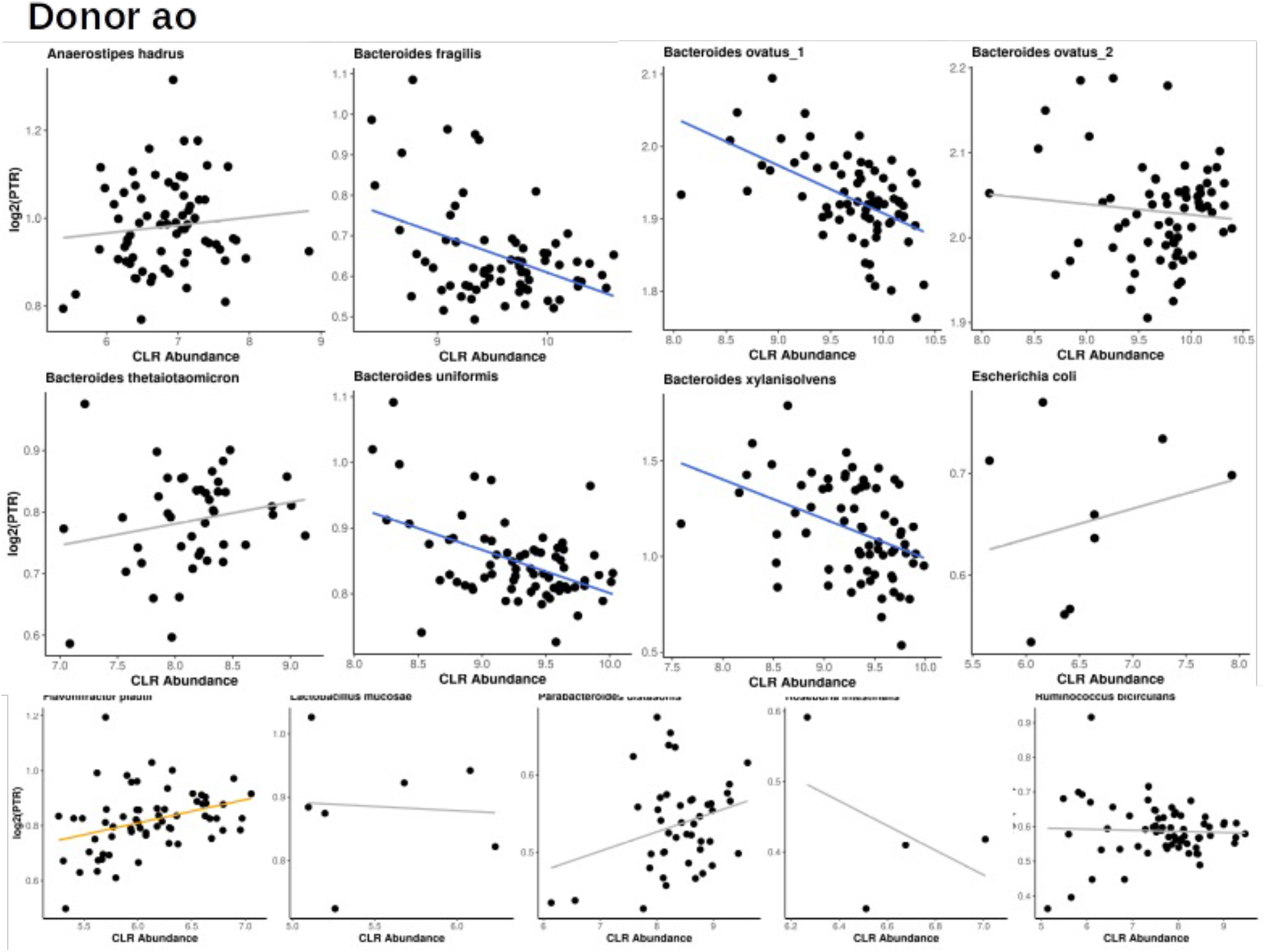
Relationships between abundance and log2(PTR) for individual taxon in donor. **ao**. Abundant taxa with relatively dense longitudinal PTR and abundance data (at least 5 matched data points; time differences between adjacent samples less than three days) were selected for analysis. Gray trend lines show no significant correlations, orange trend lines indicate significant positive correlations, and blue trend lines represent significant negative correlations (linear regression, BH-FDR < 0.05).

